# Accurate and Efficient Cell Lineage Tree Inference from Noisy Single Cell Data: the Maximum Likelihood Perfect Phylogeny Approach

**DOI:** 10.1101/742395

**Authors:** Yufeng Wu

## Abstract

Cells in an organism share a common evolutionary history, called cell lineage tree. Cell lineage tree can be inferred from single cell genotypes at genomic variation sites. Cell lineage tree inference from noisy single cell data is a challenging computational problem. Most existing methods for cell lineage tree inference assume uniform uncertainty in genotypes. A key missing aspect is that real single cell data usually has non-uniform uncertainty in individual genotypes. Moreover, existing methods are often sampling-based and can be very slow for large data.

In this paper, we propose a new method called ScisTree, which infers cell lineage tree and calls genotypes from noisy single cell genotype data. Different from most existing approaches, ScisTree works with genotype probabilities of individual genotypes (which can be computed by existing single cell genotype callers). ScisTree assumes the infinite sites model. Given uncertain genotypes with individualized probabilities, ScisTree implements a fast heuristic for inferring cell lineage tree and calling the genotypes that allow the so-called perfect phylogeny and maximize the likelihood of the genotypes. Through simulation, we show that ScisTree performs well on the accuracy of inferred trees, and is much more efficient than existing methods. The efficiency of ScisTree enables new applications including imputation of the so-called doublets.

**Availability:** The program ScisTree is available for download at: https://github.com/yufengwudcs/ScisTree

## 1 Introduction

Organisms such as human usually contain a large number of cells. For human, each individual starts from a single fertilized egg, which develops into trillions of cells in an adult. Before the era of single cell sequencing, only bulk sequencing data is available, which contains sequence data from a mixture of diverse cells in an organism. This may complicate downstream analyses such as cancer evolution (see, e.g. Gerlinger *et al.* (2012); Gundem *et al.* (2015)). Recently, single cell sequencing technology is undergoing very active development (see, e.g. Shapiro *et al.* (2013); Zong *et al.* (2012)). Single cell sequencing can generate sequences from individual cells of an organism. This new technology may have profound impact on several important biological problems.

An important problem on cells concerns the evolutionary history of cells of an organism, i.e., the cell lineage tree. Cell lineage tree is a rooted tree with extant cells at the leaves. It is highly desirable to infer cell lineage tree with single cell data (e.g., single cell DNA sequences) from multiple sampled (healthy or tumor) cells. Since phylogeny inference is a mature area in computational biology, it is tempting to use existing phylogeny inference methods that are developed for species evolution. However, cell lineage tree inference has several unique properties that make it different from traditional phylogeny inference. One of the most important aspects about single cell data is that single cell data is very noisy (see, e.g. Shapiro *et al.* (2013); Navin and Chen (2016)). While in principle genetic variations such as single nucleotide variants (SNVs) can be called from single cell data, the called SNVs tend to be very noisy due to technological errors such as allelic dropouts and genotyping errors. Existing phylogeny inference methods usually don’t explicitly consider such errors.

During the past several years, several cell lineage tree inference methods (e.g. Jahn *et al.* (2016); Ross and Markowetz (2016); Zafar *et al.* (2017)) have been developed. An assumption often made about single cell evolution is the so-called infinite sites (IS) model, which assumes there is a single mutation for one SNV. Several methods such as SCITE (Jahn *et al.*, 2016) are based on the IS model. There are also methods that don’t assume the IS model. For example, SiFit (Zafar *et al.*, 2017) assumes the finite sites model. While these methods are certainly useful, there are several major drawbacks. First, almost all existing methods (e.g., SCITE and SiFit) are based on Markov chain Monte Carlo (MCMC) and are slow for large data. With the rapid technology development, the size of single cell data is likely to increase rapidly. Many current methods don’t scale well when the number of cells or sites increases. Moreover, while existing methods do consider errors in the data, these methods usually assume uniformly distributed errors in genotypes. That is, two genotypes with the same value are assumed to have the same uncertainty no matter which cells or sites they are from. In practice, however, genotypes called from single cell sequence data tend to have “non-uniform” uncertainty. For example, a genotype with a large number of mapped sequence reads can be more reliably called than a genotype with few or even no reads. Treating genotypes uniformly may lead to loss of useful information contained in single cell sequence data. To the best of our knowledge, the only existing cell linage tree inference method that works with non-uniform uncertainty data is SCIΦ (Singer *et al.*, 2018). SCIΦ works with mapped sequence reads from single cells and calls genotypes and infers cell lineage tree simultaneously. While SCIΦ is attractive in principle, a main downside of SCIΦ is that it is also based on MCMC and is slow for large data. For single cell genotype calling from single cell sequence reads, Monovar (Zafar *et al.*, 2016) is currently the commonly used approach.

In this paper, we develop a new method called ScisTree (which stands for {S}ingle {c} ell {i}nfinite {s}ites {Tree}) for inferring cell lineage tree and calling genotypes simultaneously from single cell data with non-uniform uncertainty. ScisTree assumes the infinite sites model. The following are the features of the ScisTree approach.

1. ScisTree takes uncertain single cell genotypes as input. Different from most existing methods (e.g., SCITE and SiFit), each genotype (i.e., the genetic state of a cell at a SNV site) can have its own individualized probability (called genotype probability). This allows more realistic representation of uncertain genotypes, where the content of single cell sequence data (e.g., sequence depth) may vary at different sites and cells. Different from SCIΦ, ScisTree doesn’t work with single cell sequence reads directly. Instead, ScisTree assumes that genotype probabilities are estimated from single cell sequence reads first. This is indeed feasible: for example, Monovar outputs estimated genotype probability for each called genotype. We have also developed our own approach for calculating genotype probability from sequence reads. Simulation shows that ScisTree outperforms existing methods which assume uniform genotype uncertainty when genotype uncertainty is unevenly distributed in the data. Representing uncertain genotypes by individualized genotype probabilities also provides a natural means to address the missing value issue. For example, for missing data without prior knowledge in the binary genotype case, we can simply set the probability of both 0 and 1 genotypes to 0.5.
2. ScisTree is a maximum likelihood approach. Genotypes are called by choosing genotypes that maximize the likelihood. While finding the maximum likelihood cell lineage tree is intractable in general, ScisTree implements a fast heuristic for finding the cell lineage tree that maximizes the genotype probability under the infinite sites model over the tree space. The tree space search is based on an efficiently computable likelihood. This makes ScisTree efficient and scalable to hundreds of cells and thousands of SNV sites. Simulation shows that ScisTree is much faster than existing methods while its cell lineage tree inference accuracy is better than or similar to those methods.
3. ScisTree allows both binary and ternary genotypes. ScisTree can also address various single cell data noises such as doublets. Sequence reads from a doublet are from more than one cell due to noises in data collection. ScisTree can impute doublets from the given uncertain genotypes. To the best of our knowledge, ScisTree is the first method for imputing doublets. Note that there are existing methods (e.g., SCITE) that can be configured to work with data with doublets; but these methods don’t attempt to impute the doublets.

The program ScisTree is available for download at: https://github.com/yufengwudcs/ScisTree.

## 2 Background

ScisTree assumes the infinite sites (IS) model. The IS model assumes that a mutation in the evolutionary history can occur at most once at a single site. Under the IS model, all (and only the) cells with the mutant allele at a SNV site are clustered in a single subtree of the cell lineage tree. ScisTree takes uncertain genotypes at SNV sites as input. Genotype considered in this paper is either binary or ternary. We let *G* be a genotype matrix of *n* rows by *m* columns, where *n* is the number of cells and *m* is the number of SNV sites. For uncertain genotypes, genotype probability is defined by the probability function 𝒫. For each genotype *G*[*i, j*] for cell *i* and site *j*, we let 𝒫_*g,i,j*_ be the probability of *G*[*i, j*] = *g* for the genotype state *g*. For binary genotypes, *g* ∈ {0, 1} where 0 refers to the wild-type and 1 refers to the mutant. 𝒫_0,*i,j*_ + 𝒫_1, *i, j*_ = 1.0 for all *i* and *j*, where 1 ≤ *i* ≤ *n* and 1 ≤ *j* ≤ *m*. For ternary genotypes, *g ∈* {0, 1, 2} where 0 is the homozygous wild-type, 1 is the heterozygote and 2 is the homozygous mutant. 𝒫_0,*i,j*_ + 𝒫_1,*i,j*_ + 𝒫_2,*i,j*_ = 1.0 for all *i* and *j*. Note that existing single cell genotype calling tools (e.g., Zafar *et al.* (2016)) output (ternary) genotype probabilities, which can be converted to binary genotype probabilities (i.e., wild-type and mutant). We define the genotype probability for a fixed genotype matrix *G* to be 𝒫(*G*) Π = _*i,j*_𝒫_*G*[*i,j*],*i,j*_. We assume all-0 genotypes at the root of cell lineage tree. For the ease of exposition, in the following, we use binary genotypes by default, unless otherwise stated. Note that if the IS model holds for each site, only binary genotypes will be observed. In practice, however, homozygous mutants may be observed due to more complex evolutionary processes, such as recurrent mutations and copy number changes. If such complex processes occur at a site, the IS model may not hold for the site. While the IS model has been popular in the literature on single cell evolution, there are also papers that argue the IS model should be scrutinized (Kuipers *et al.*, 2017; Zafar *et al.*, 2017). In this paper, ScisTree assumes such complex processes, if present, tend to be rare. Under this assumption, a majority of sites are likely to support the IS model.

When there is no uncertainty in genotypes and the number of SNV sites is large, inferring cell lineage tree is straightforward. Cell lineage tree inference from fixed binary genotypes under the IS model is essentially the classic (binary) perfect phylogeny problem. There exists a well-known linear time algorithm by Gusfield (Gusfield, 1991) for this problem. The main difficulty for cell lineage tree inference is the inherent complexity in single cell data. There are two categories of complexity: evolutionary complexity and technological complexity (see, e.g., Shapiro *et al.* (2013); Navin and Chen (2016)).

### Evolutionary complexity

Evolutionary complexity for single cells refers to complex evolutionary processes in single cells. For example:

1. Recurrent mutation. There is more than one mutation at a site.
2. Copy number changes. This may occur in any cell, especially a tumor cell. Copy number changes may involve copy number gain or loss. Copy number loss may lead to loss of heterozygosity (LOH).

### Technological complexity

There are technical difficulty of obtaining accurate single cell sequence data. For example,

1. Allelic dropout (or simply dropout). A segment of genomic region may be missing from sampled DNA prior to sequencing. Dropout may lead to errors in genotype calling. For example, consider a heterozygous genotype. If dropout occurs at the mutant allele, the genotype may be mistaken for a homozygous wild-type.
2. Reads error. The allele contained in a mapped sequence read may be different from the true allele. The probability of error in reads is usually much smaller than the dropout rate but is not zero.
3. Doublets. Single cell sequencing usually needs to separate cells into single cells. This process is not perfect. Sometimes, two or more cells may be mistaken to be a single cell.
4. Sequence coverage. Single cell sequence data may have low coverage: there may be few or even no sequence reads at some SNV sites. This leads to more uncertain genotype calling.

In this paper, we focus on addressing technological complexity in single cell data. In particular, we use uncertain genotypes with individualized genotype probabilities to accommodate technological noises. For evolutionary complexity, although we don’t explicitly address these issues, our model is flexible enough to at least tolerate some of these complexities. We will show that our method is still reasonably accurate when the deviation from the assumed IS model is modest.

## 3 Methods

### 3.1 The high-level approach

Under the infinite sites model, fixed genotypes uniquely determine a (possibly multifurcating) phylogeny (see, e.g., Gusfield (1997, 2014)). When the number of SNV sites increases, this phylogeny (called perfect phylogeny) is increasingly more likely to be the true cell lineage tree topology. Thus, with enough sites, the maximum likelihood cell lineage tree topology is exactly the binary perfect phylogeny under the IS model. It can be shown that when uniform prior is assumed for *G* and *T*, we can use 𝒫(*G*) as the likelihood function, with the condition that *G* satisfies the IS model. That is, we want to find the genotypes *G* that satisfy the IS model (and thus allow a perfect phylogeny, which is used as the inferred cell lineage tree) and 𝒫(*G*) is maximized. To be more specific:

#### Maximum likelihood perfect phylogeny problem

given the genotype probability 𝒫 of each genotype for *n* cells and *m* sites, find the cell lineage tree topology *T*_*opt*_ and the genotype matrix *G*_*opt*_ such that *G*_*opt*_ satisfies the IS model (and thus *T*_*opt*_ can be constructed from *G*_*opt*_ via the perfect phylogeny formulation) and 𝒫(*G*_*opt*_) is maximized.

Figure 1 gives an illustration of the maximum likelihood perfect phylogeny problem. The true single cell phylogeny is shown in Figure 1(a). The single cell genotypes and sequence reads are shown in Figure 1(b). Here dropout can occur at some alleles, where there are no reads. Sequencing error may also occur. Genotypes called from the sequence reads are shown in Figure 1(d). Due to dropouts and read errors, the called genotypes are different in several positions from the true genotypes in Figure 1(c). In Figure 1(d), there are two wrongly called genotypes (in bold face) due to dropout. Uncertain genotypes with uniform probability (of genotype 0) are shown in Figure 1(e). The true genotypes would lead to maximum probability for the uniform probability shown in Figure 1(e). However, an equally optimal solution is shown in Figure 1(f). This indicates that uniform probability is not informative enough for distinguishing these two solutions. Non-uniform genotype probability can be more realistic for sequence data. Figure 1(g) shows a possible setting for non-uniform genotype probability. For example, at site *S*_1_ and cell *C*_1_, the probability of being genotype 0 is reduced to 0.7 because the number of reads with allele-0 is less than expected. That is, it is likely that a dropout occurs at this position. This is different from the genotype of *C*_1_ at *S*_4_, where only allele-0 reads exist and the number of reads is more than expected. So *G*[*C*_1_, *S*_1_] is more likely to be of genotype 1 than *G*[*C*_1_, *S*_4_]. In order to obtain genotypes satisfying the IS model, the optimal genotypes have *G*[*C*_1_, *S*_1_] changed to 1 and *G*[*C*_5_, *S*_3_] changed to 0 from the originally called genotypes. Note that other ways to make the genotypes satisfying the IS model would lead to lower probability. One such example is the genotypes in Figure 1(f) (where *G*[*C*_3_, *S*_2_] is set to 1 instead of *G*[*C*_1_, *S*_1_] based on genotypes in Figure 1(d), and *G*[*C*_3_, *S*_3_] is set to 0 instead of *G*[*C*_5_, *S*_3_]): 𝒫_1,*C*3_,*S*_2_ = 0.05 *<* 𝒫_1,*C*1_,*S*_1_ = 0.3; also, 𝒫_0,*C*3_,*S*_3_ = 0.01 *<* 𝒫_0,*C*5_,*S*_3_ = 0.02.

**Figure 1:**
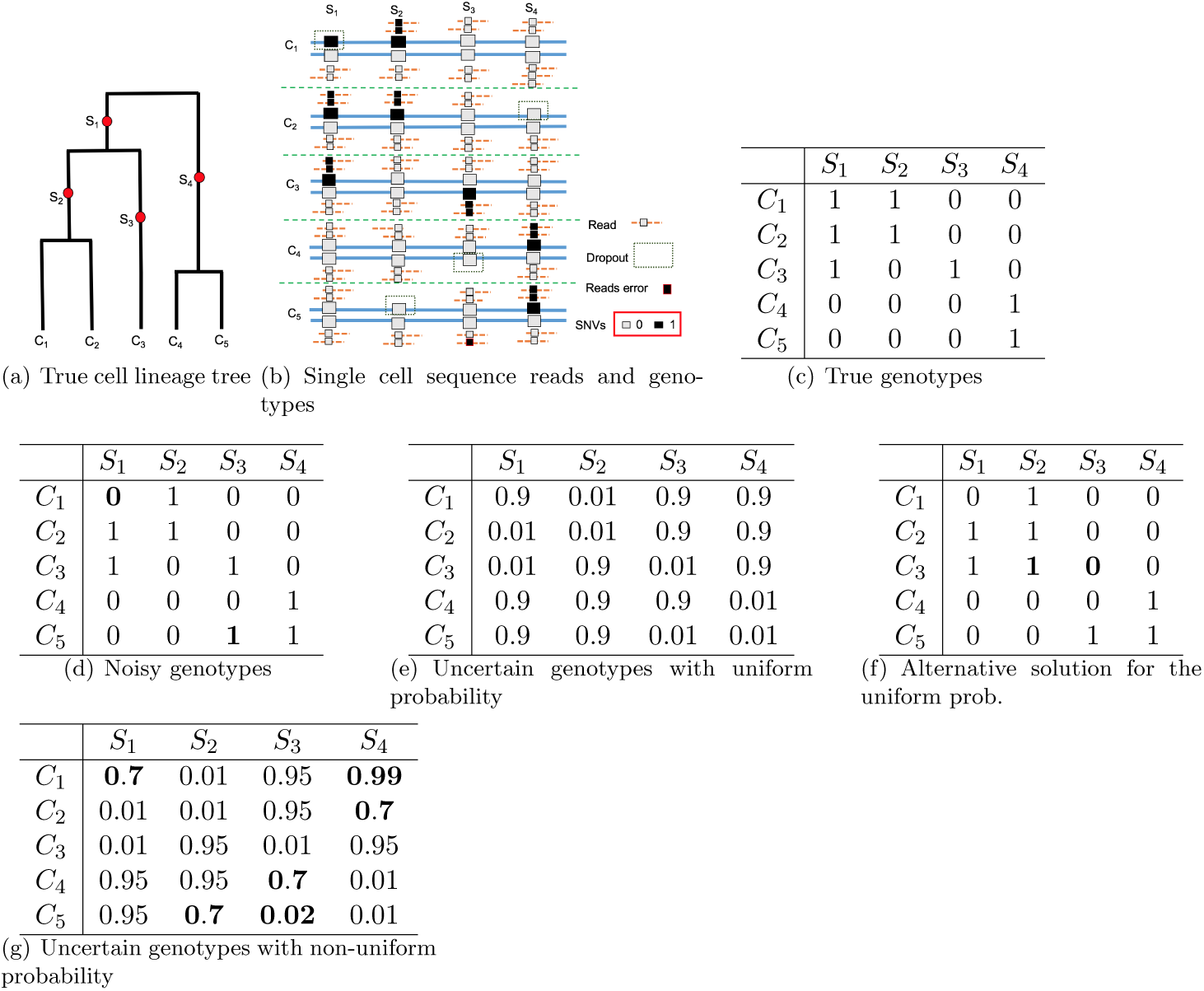
An illustration of cell lineage tree and uncertain genotypes. Part 1(a): the true cell lineage tree with five cells and four sites (with mutations labeling the branches). Part 1(b): sequence reads (lines with smaller boxes) and single cell genotypes (larger boxes) at the four SNV sites. Read depth may vary. Dropout and reads errors may occur. Part 1(c): true (binary) genotypes. Part 1(d): genotypes called from reads. Part 1(e): uncertain genotypes with uniform probability model (assuming same genotype state with identical probability). The probability shown in the table is for the genotype state 0. In the example, genotype state 0 has probability 0.9 and genotype state 1 has probability 0.99 (and thus 0.01 of being genotype state 0). Part 1(f) shows an alternative solution to maximum likelihood perfect phylogeny problem for the uniform probability given in Part 1(e). Positions changed from the noisy genotypes are in bold face. Part 1(g): uncertain genotypes with non-uniform probability. Positions with more or less than expected number of reads are in bold face.

The maximum likelihood perfect phylogeny problem is intractable in general (see Section 3.2). To develop a practical method, we adopt a heuristic algorithm that is implemented in a software tool called ScisTree. ScisTree takes the following iterative approach:

1. Construct an initial rooted binary tree *T*_0_ based on 𝒫. This is done by finding genotypes *G*_0_ that maximize 𝒫(*G*_0_*|T*_0_). Initialize *T*_*opt*_ *← T*_0_, 𝒫_*opt*_ *←*𝒫(*G*_0_*|T*) and *G*_*opt*_ *← G*_0_.
2. Find rooted binary trees 𝒯_*c*_ that are similar topologically to *T*_*opt*_.
3. Let *T ∈𝒯*_*c*_ that maximizes the likelihood 𝒫(*G|T*) for some genotypes *G*. If 𝒫(*G|T*) *> 𝒫*_*opt*_, set *T*_*opt*_ *← T*, 𝒫_*opt*_ *←𝒫*(*G|T*), *G*_*opt*_ *← G* and go to step 2. Otherwise, stop.

Note that 𝒫(*G|T*) is the genotype probability for genotype *G* where *G* is consistent with tree *T* under the IS model. The key idea is relying the underlying tree *T* to find *G* that maximizes 𝒫(*G*). When the procedure finishes, we obtain both the optimal (or near-optimal) rooted binary cell lineage tree *T*_*opt*_ and the genotypes *G*_*opt*_ that (approximately) maximize the genotype probability. The key step in the above procedure is computing the maximum probability of uncertain genotypes under the genotype probability 𝒫 for a fixed phylogeny *T*, and also finding such optimal genotypes. We show that in the following, finding *G* that maximizes 𝒫(*G|T*) for a given *T* has a linear-time algorithm.

### 3.2 Complexity of the maximum likelihood perfect phylogeny problem

Theorem 3.1 shows that the maximum likelihood perfect phylogeny problem is a challenging computational problem. Due to the space limit, its proof is given in the Supplemental Materials.

#### Theorem 3.1

*The maximum likelihood perfect phylogeny problem is NP complete.*

### 3.3 Finding genotypes *G* that maximize its probability for a fixed tree

Suppose we are given a rooted binary tree *T*. We want to find genotypes *G* that satisfy the IS model and maximize 𝒫(*G|T*) where *T* is the underlying cell lineage tree. We denote *𝒫*(*G|T*) as the maximum likelihood of genotypes for *T*. We first consider the binary genotype case. Recall that under the IS model, the set of mutant genotypes at a site *s* correspond to a subtree in *T* and the single mutation occurs on the branch out of the subtree root. We define 𝒫_*s,v*_(*G|T*) for a node *v* and a site *s* to be the probability of all genotypes at site *s* given that the mutation at *s* occurs on the branch out of this subtree that is rooted at *v*. Since *T* is fixed, we can examine each subtree rooted at node *v* and compute 𝒫_*s,v*_(*G|T*). The maximum probability at site *s* is:

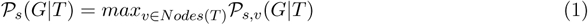

Here, *Nodes*(*T*) is the set of nodes in *T*. The probability of genotypes is simply the product of the probability of each site. More precisely, for each node *v* in *T* :

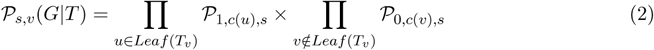

Here *T*_*v*_ refers to the subtree rooted at node *v*. *Leaf* (*T*_*v*_) is the set of leaves of *T*_*v*_. And *c*(*u*) is the cell corresponding to a leaf *u* in *T*.

Equations 1 and 2 lead to a simple algorithm for computing the maximum probability 𝒫_*s*_(*G|T*) for each site *s*. Computing 𝒫_*s,v*_(*G|T*) for each *v* directly would lead to *O*(*n*^2^) time for each site. A simple observation is that we can apply dynamic programming by taking a bottom-up approach as follows. We define *q*_*v*_ as the *ratio* of the probability of the genotypes within the subtree *T*_*v*_ being genotype 1 and the probability of these genotypes being genotype 0. We have:

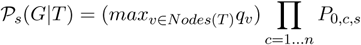

The algorithm for computing 𝒫_*s*_(*G|T*) based on *q*_*v*_ for a single site *s* for a fixed binary tree *T* is given below, which has the running time of *O*(*n*). Computing 𝒫(*G|T*) involves calculating the product of 𝒫_*s*_(*G|T*) over each of *m* sites and thus takes *O*(*mn*) time.

#### Algorithm 1 Maximum probability computation of binary genotypes

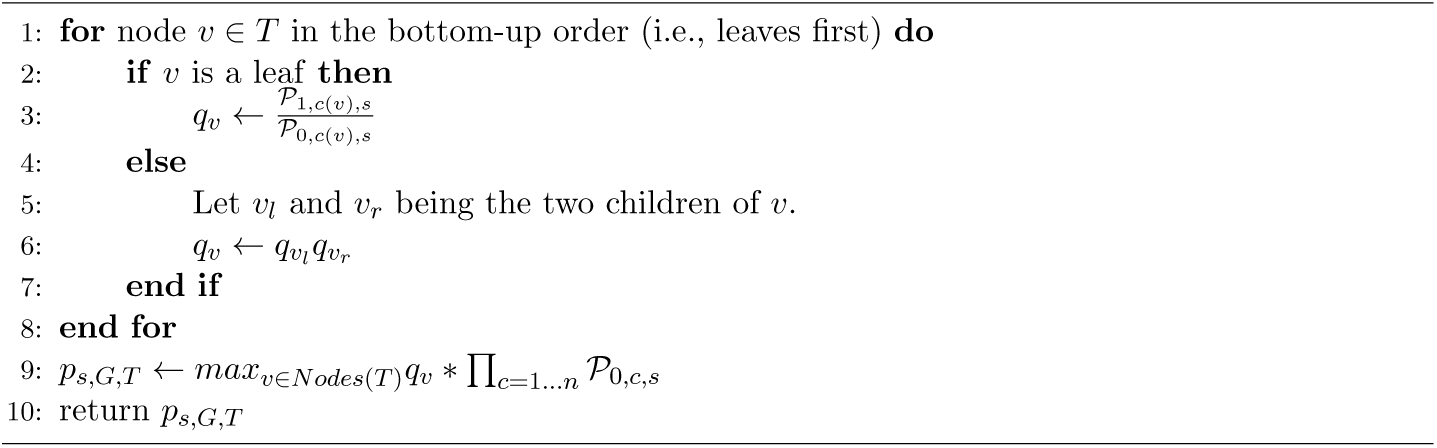

#### Genotype calling

Note that the optimal genotypes at site *s* corresponding to 𝒫_*s*_(*G|T*) can be easily found in the algorithm by keeping track of what subtree root *v*_*opt*_ gives the maximum genotype probability. That is, at site *s*, cells within the subtree rooted at *v*_*opt*_ have the mutant genotype 1, while other cells have the wild-type genotype 0.

#### The ternary genotype case

Algorithm 1 can be extended to the case of ternary genotypes. The details are given in the Supplemental Materials.

### 3.4 Finding optimal cell lineage tree

For a given cell lineage tree *T*, we can compute the maximum probability of genotypes using Algorithm 1. Since the space of trees is immense, it is infeasible to examine every possible *T*. ScisTree takes the following heuristic for searching the tree space to find the optimal cell lineage tree.

#### 3.4.1 Initial tree construction

ScisTree constructs an initial cell lineage tree as follows. It first constructs a fixed genotype matrix *G*_0_ from the given uncertain genotypes. *G*_0_ is obtained by taking the most probable genotype at each position in the matrix. We break ties arbitrarily. This *G*_0_ is called the maximal probability genotype matrix. We then use the well-known neighbor joining method (Saitou and Nei, 1987) to construct the initial tree from *G*_0_. To apply neighbor joining, we define the distance *d*(*i, j*) between two cells *i* and *j* as follows. For binary genotypes, *d*(*i, j*) is simply the Hamming distance between the two genotype vectors for cells *i* and *j* in *G*_0_. For ternary genotypes, we use the summation of absolute allele differences between two cells *i* and *j* to be the distance: 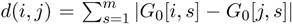. By default, ScisTree uses all genotypes when calculating the distance even when some genotypes have significant uncertainty. Our experience indicates that this may lead to poor initial trees given data with significant noise. Thus, ScisTree can be configured to only use more certain genotypes for distance calculation. In particular, ScisTree has a threshold value *t*_*d*_ (default to be zero) which is the minimum difference between the probabilities of different genotypic states. For example, if *t*_*d*_ = 0.9 for the case of binary genotypes, only genotypes with probability for genotype 0 is no less than 0.95 or no greater than 0.05 are used in distance calculation. Note that this is another benefit of using individualized genotype probabilities. We use all-0 genotypes as the outgroup to root the tree constructed by neighbor joining. The resulting initial tree is thus a rooted binary tree.

#### 3.4.2 Tree space search

Starting from the constructed initial tree, ScisTree searches for the optimal tree topology by exploring the neighborhood of the current best tree. In particular, ScisTree examines the trees found by a single nearest-neighbor interchange (NNI) from the current best tree. We choose NNI because it is more scalable than other types of tree neighborhood search (e.g., subtree prune and regraft). Maximum genotype probability is computed for each candidate tree found by NNI. If a candidate tree has higher genotype probability than the current best tree, the candidate tree is adopted to be the new best tree. The search stops when none of the neighboring trees has higher likelihood than the current best tree. Note that the number of NNI trees for the current tree is *O*(*n*). So the total running time is *O*(*mn*^2^*N*), where *N* is the number of iterations in search.

### 3.5 Doublet imputation

One main source of technological complexity in single cell data is doublet. In single cell genotype data, some genotypes that are supposed to be from a single cell can be from a mixture of two or more cells. Such a cell is called doublet. Since doublets may lead to artifacts in downstream analyses (e.g. cell lineage tree inference or genotype calling), it is desirable to impute doublets from the given uncertain genotypes.

Thanks for the efficiency of the maximum likelihood approach, we have extended the ScisTree approach to impute doublets from noisy single cell data. For simplicity, in the following, we assume the number of doublets *n*_*d*_ is provided by the user. In practice, the exact number of doublets may not be known. In general, we note that while some doublets are relatively obvious to impute, it is difficult to impute all doublets. When the specified doublet number is larger than the number of “obvious” doublets, ScisTree tends to stop reporting meaningful doublets: it simply reports trivial doublets (i.e., doublets that are formed by the same cells). ScisTree stops the imputation when such trivial doublets are imputed. Thus, the user can choose a large value as the upper bound for the number of doublets, if the number of true doublets is not known.

ScisTree takes the following iterative algorithm for imputing doublets.

#### Algorithm 2 An iterative approach for doublet imputation

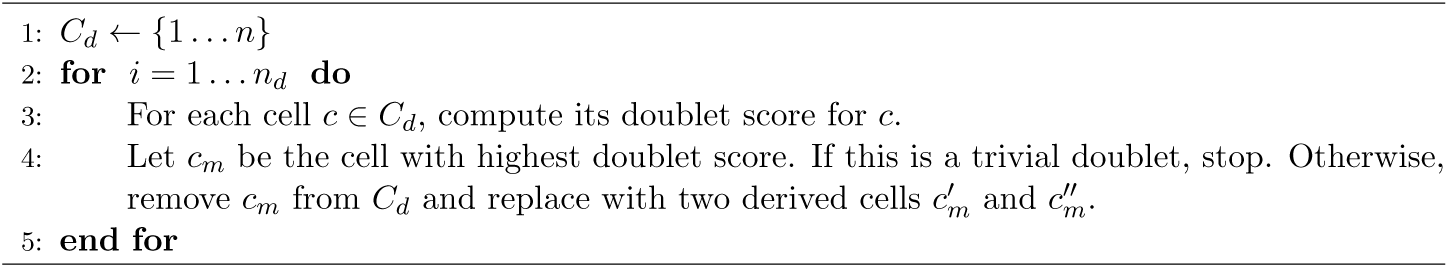

The key step in Algorithm 2 is scoring a candidate cell as follows. ScisTree first calls genotypes of all other cells except the doublet candidate using the maximum likelihood perfect phylogeny approach in Section 3.1; then it finds the most probable splitting of the candidate cell into two derived cells such that genotypes of the two derived cells are still consistent with the IS model; the candidate score is the maximum probability of splitting the candidate cell. Once the optimal score is obtained, genotypes can be called for the two new cells by optimally splitting a doublet cell into two derived cells. Overall, doublet imputation takes *O*(*n*_*d*_*n*^2^*m*(*m* + *n*)*N*) = *O*(*n*^3^*m*(*m* + *n*)*N*) time, where *N* is the number of iterations for finding the maximum likelihood tree. Due to the space limit, the algorithm is given in the Supplemental Materials.

### 3.6 Read counts simulation

We assume sequence depth follows a normal distribution at each site by default, unless otherwise stated. For simplicity, at the same site, different cells are assumed to have the same expected sequence depth. We let *µ*_*d*_ be the average read depth, and *σ*_*d*_ be the standard deviation of the normal distribution of read depth. We simulate read counts for each genotype at each site. Each copy (allele) of the genotype is sampled with equal probability. If a dropout occurs at one of the two copies, all the reads from that copy are discarded.

### 3.7 Genotype probability computation

Given mapped single cell sequence reads, existing single cell genotype calling tools (e.g., Monovar) can compute genotype probability for each called genotype. In this paper, we compute genotype probabilities from allele read counts in the following way (similar to the approach in Duitama *et al.* (2011)). Our experience suggests that genotype probabilities computed this way perform slightly better than those computed by Monovar in our simulation. Let *R* = {*r*_0_, *r*_1_} be a set of reads sampled for a cell *c* at a site *s*. Here, *r*_0_ (respectively *r*_1_) is the number of reads with allele 0 (respectively 1) at the site. Note that the number of reads at a site depends on whether dropout occurs, and dropout tends to reduce the number of reads. We first estimate the read depth distribution *P*_*s*_(*k*) for site *s*, assuming normal distribution. Here, *P*_*s*_(*k*) is the probability of having *k* reads for one allele at site *s*. If both alleles of a genotype have dropout, this leads to a missing genotype. For the three possible genotypes *g ∈* {0, 1, 2}, we calculate the posterior probability of *g* given *R* as follows. For *g* = 0,

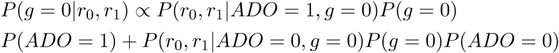

Here, *ADO* is a binary random variable, where *ADO* = 1 indicates an allelic dropout occurs, and *ADO* = 0 indicates no dropout. We have 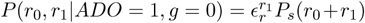, where *ϵ*_*r*_ is the reads error rate (which is assumed to be known). Also, 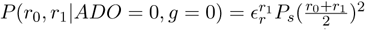, where we assume reads are evenly distributed between two genotype alleles. *P* (*g* = 1*|r*_0_, *r*_1_) and *P* (*g* = 2*|r*_0_, *r*_1_) can be calculated in the similar way. The prior probability *P* (*g*) is calculated using the estimated allele frequency, assuming the Hardy-Weinberg equilibrium. The prior probability for dropouts *P* (*ADO* = 1) can be assigned uniformly to the global dropout rate.

## 4 Results

### 4.1 Cell lineage tree inference from simulated genotypes

Table 1 lists the simulation parameters and their default values. By default, we simulate binary genotypes. The default number of sites *m* is either five or ten times of the number of cells *n*. For each setting, we simulate 50 replicates. Reported results are the average over these replicates.

**Table 1:** A list of parameters and their default values used in the simulation.

| Description | Symbol | Default |
| --- | --- | --- |
| The number of cells | $n$ | 100 |
| The number of SNV sites | $m$ | 500/1,000 |
| Deviation from clock property in true cell lineage tree | $t_0$ | 0.1 |
| Recurrent mutation rate | $r_m$ | 0.0 |
| Copy number loss rate | $m_c$ | 0.0 |
| Genotype error rate | $\epsilon_g$ | 0.002 |
| Allelic dropout rate | $\epsilon_d$ | 0.2 |
| Average read depth of a single allele | $\mu_d$ | 4 |
| Standard deviation of the read depth | $\sigma_d$ | $\frac{1}{2}\mu_d$ |

For cell lineage tree inference, we first compare ScisTree with two methods, SCITE (Jahn *et al.*, 2016) and SiFit (Zafar *et al.*, 2017). We choose SCITE because a previous comparison study (Miura *et al.*, 2018) suggests SCITE performs well when compared with other existing methods. We choose SiFit because it doesn’t assume the IS model; comparing with it may allow evaluation of the effect of modeling assumptions. Since cell lineage tree can be inferred using classic phylogeny inference methods, we also compare with neighbor joining (NJ). Both SCITE and SiFit are run with true false negative and false positive rates. We run 90,000 iterations for SCITE, and run 200,000 iterations for SiFit, unless otherwise stated. Note that the choices on the number of iterations here may affect accuracy, although it is difficult to pick the best number.

We use the well-known Robinson-Foulds (RF) distance between the inferred cell lineage tree *T* and the true tree *T′* as the main measure of phylogenetic accuracy. The RF distance is equal to the number of clades that are in *T* but not in *T′*. We normalize the RF distance to be between 0 and 1 (i.e., RF distance divided by *n -* 2, the maximum number of non-singleton clades in a rooted tree with *n* leaves). See Section 4.1.1 for alternative metrics for benchmarking. To test the performance of cell lineage tree inference, we vary a number of parameters, including *n, m, t*_0_, *r*_*m*_, *m*_*c*_, *ϵ*_*g*_ and *ϵ*_*d*_ (see Table 1).

We now evaluate the performance of ScisTree with *non-uniform* genotype probability data (calculated using the procedure described in Section 3.7). To test the performance of ScisTree with more sites, we let *n* = 100 and *m* = 1, 000. Since both SCITE and SiFit don’t consider non-uniform genotype probability and only work with uniform error rates, we use the maximal probability genotypes as the called genotypes, along with the true false positive and false negative rates when running SCITE and SiFit (and neighbor joining). To see the difference between the uniform and non-uniform probability settings, we also run ScisTree with the uniform genotype probability as computed from the maximal probability genotypes under the true false positive and negative rates.

The results are shown in Figure 2. It can be seen that ScisTree with non-uniform genotype probability clearly outperforms SiFit, ScisTree with uniform probability and neighbor joining. SCITE still performs well but now becomes slightly less accurate than ScisTree in most cases. When the read depth is low (say at 2x), SCITE is slightly more accurate than ScisTree with non-uniform probability. As shown in Figure 2(a), increasing read depth (while setting read depth standard deviation to be half of the read depth) appears to let ScisTree significantly outperform other methods (including SCITE). Note that SCITE and other methods only improve accuracy slightly (or even decrease sometimes) when read depth increases. This may be because that when read depth is high, probabilities at different genotypes become more non-uniform (note that standard deviation also increase, since standard deviation is fixed to be half of the average); assuming uniform error rates may be less realistic in this case. For comparison, in the uniform probability case, SCITE can be slightly more accurate than ScisTree in some cases. This suggests that using non-uniform genotype probability can indeed improve the tree inference accuracy.

**Figure 2:**
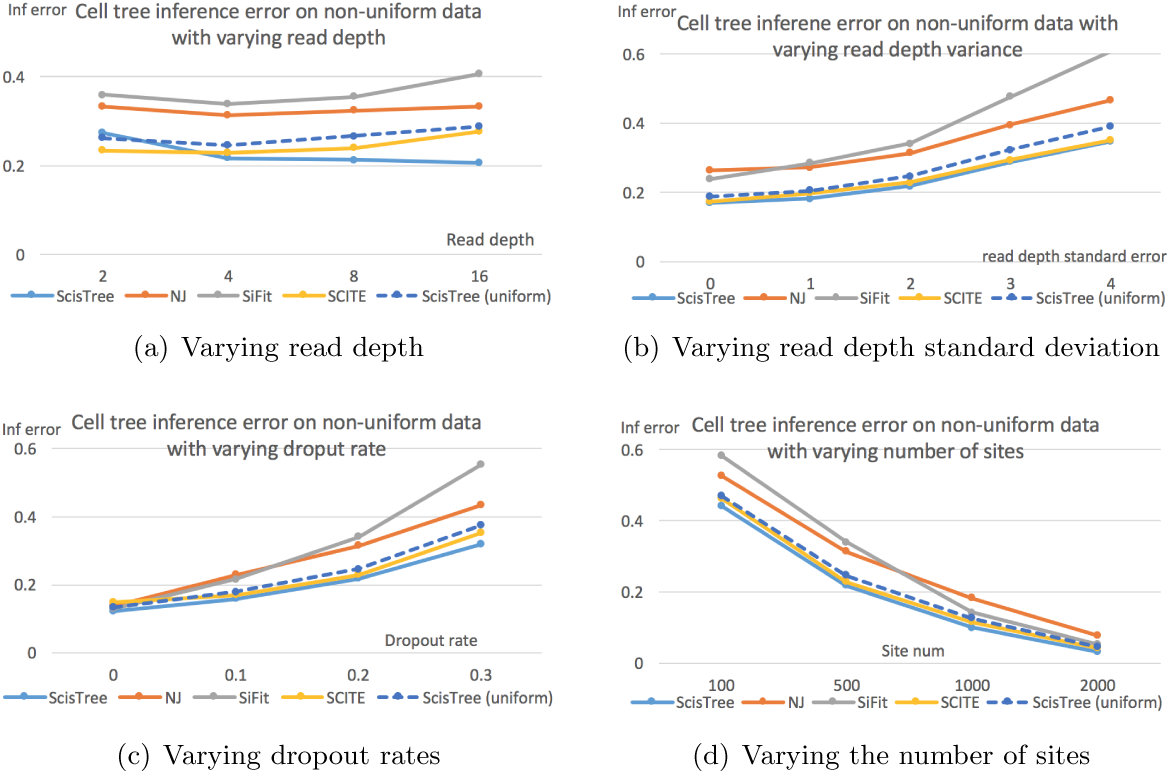
Cell lineage tree inference error by ScisTree with non-uniform probability computed by sequence reads, and neighbor joining (NJ) and ScisTree on uniform probability. Part 2(a) shows the effect of increasing mean read depth (with read depth standard deviation being half of the mean read depth). Part 2(b) shows the effect of increasing read depth standard deviation. Part 2(c) shows the effect of increasing dropout rates. Part 2(d) shows the effect of number of sites. Y axis: inference accuracy (error) in terms of average RF distance between true tree and inferred tree. Default settings of parameters are used for parameters not being tested.

#### 4.1.1 Clade accuracy for cell lineage tree inference

There are other types of phylogenetic accuracy measures (other than the RF distance) for benchmarking the cell lineage tree inference. Here, we consider the following two measures (which are similar to those in the Appendix of the preprint: “Integrative inference of subclonal tumour evolution from single-cell and bulk sequencing data” by S. Malikic, et al). These two measures work with mutation trees (instead of cell lineage trees). We suggest the readers to read the literature on tumor phylogeny inference for more detailed description of mutation (or clonal) trees. Briefly, mutation tree focuses on the timing order of mutations (i.e., site labels) on the cell tree. We say a mutation *m*_*a*_ is ancestral to *m*_*b*_ if the clade formed by cells carrying mutant *m*_*a*_ contains the clade formed by cells carrying mutant *m*_*b*_.

1. Ancestor-descendant error: the percentage of pairs of mutations *m*_*a*_ and *m*_*b*_ where *m*_*a*_ is ancestral to *m*_*b*_ in the true tree but *not* so in the inferred tree. Note: we use “error” to be consistent with the Robinson-Foulds distance (which measures errors).
2. Different-lineage error: the percentage of pairs of mutations *m*_*a*_ and *m*_*b*_ where *m*_*a*_ is *not* ancestral to *m*_*b*_ in the true tree but *m*_*a*_ is ancestral to *m*_*b*_ in the inferred tree.

We simulate 100 cells with various numbers of SNV sites. For comparison, we compare ScisTree with SCITE under these two measures of inference errors. Results are shown in Figure 3. Our results show that ScisTree appears to outperform SCITE significantly under these two error measures on mutation trees. Our experience indicates that SCITE tends to infer mutation tree with long nested mutations that are not in the original simulated mutation trees. This may explain why SCITE doesn’t perform well under these two measures.

**Figure 3:**
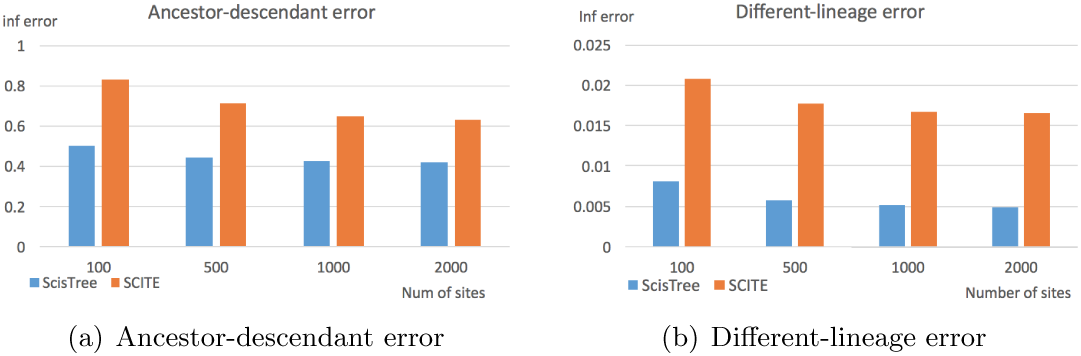
Mutation tree inference error (the ancestor-descendant error and the different lineage error) by ScisTree and SCITE. 100 cells. Number of sites: from 100 to 2,000. Y axis: error measure of mutation tree between true mutation tree and inferred mutation tree.

### 4.2 Cell lineage tree inference from simulated sequence reads

So far we use simulated genotype read counts without simulating sequence reads. In practice, current single cell data is in the form of sequence reads. We now simulate single cell sequence reads as follows. We use a segment of human chromosome 20 as the reference genome. We create alternative genomes from the simulated SNV alleles for each cell. We use the reads simulator wgsim to simulate sequence reads from the alternative genomes from the cells. We simulate four different reads coverages (for a single chromosome copy): 2, 4, 8 and 16. We then use Monovar to call single nucleotide variants (SNVs) from the reads. We then calculate the genotype probabilities based on the allelic reads counts from the VCF files by Monovar, using the approach in Section 3.7. There are some genotype positions where Monovar reports to be missing. We assign genotype probability of 0.5/0.5 for these positions. For comparison, we run SCIΦ with the simulated sequence reads (in BAM format). The suggested parameters (by its user manual) are used when running SCIΦ. We also run SCITE on the maximal probability genotypes as before.

As shown in Figure 4, ScisTree under the default settings is less accurate than SCIΦ (and SCITE) at low coverage. However, when we discard highly uncertain genotypes from initial tree construction (see Section 3.4.1), ScisTree performs significantly better than under the default settings. This can be seen by setting the probability difference threshold *t*_*d*_ to 0.95 (smaller *t*_*d*_ values also give somewhat smaller gains). Here, ScisTree clearly outperforms SCITE and has similar accuracy as SCIΦ. This suggests that accurate initial tree construction can be important for noisy data. Note that this simulation uses simulated single cell data with more noise than those in Section 4.1. This is because noise can be introduced during reads mapping and SNV calling. On the other hand, the simulated reads distribution here is different from the results shown in Figure 2(a). Our results show that dealing with non-uniform genotype uncertainty can be important for data with significant noise.

**Figure 4:**
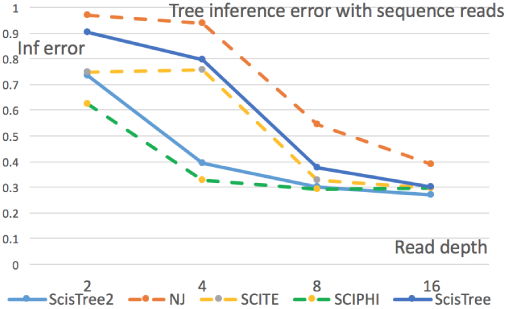
Tree inference error with simulated single cell sequence reads. Results for ScisTree (default settings), ScisTree2 (using more reliable genotypes for initial tree construction, with probability difference threshold t_d_ = 0.95), SCITE and SCIΦ and neighbor joining are reported. X axis: average reads coverage. Y axis: normalized tree inference error.

#### 4.2.1 Impact of sequence reads distribution

In most of our simulation, we assume that single cell sequence reads follow the normal distribution. In reality, single cell sequence reads may follow more complex distributions. Here, we conduct further simulation to test the effect of sequence reads distribution on the inference accuracy. In particular, we assume single cell sequences follows the beta-binomial distribution. The beta-binomial distribution has two parameters *α* and *β*, which determine the shape of the distribution. We test two settings: *α* = *β* = 2 and *α* = *β* = 4. The cell lineage tree inference error with these two settings and the case of normal distribution is shown in Figure 5. We can see that under beta-binomial distribution, cell tree inference accuracy is indeed less than the case of normal distribution. This is likely due to the overdispersion in the beta-binomial distribution. In this case, it is more difficult to get a good estimate of genotype probability than the normal distribution case. This indicates that obtaining good estimates of genotype probability with complex single cell data is likely to be an important research problem for cell lineage tree inference.

**Figure 5:**
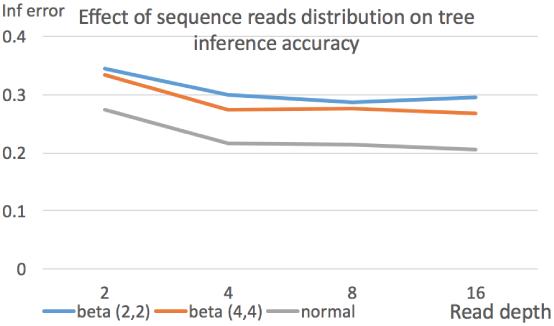
Effect of sequence reads distribution on cell lineage tree inference error. Three settings: beta-binomial (with parameters 2 and 2), beta-binomial (with parameters 4 and 4), and normal distribution. 100 cells and 1,000 sites. Average reads depth is 4.

### 4.3 Running time

Figure 6 shows the average running time for a single dataset of ScisTree, neighbor joining, SCITE and SiFit under different settings with simulated data. ScisTree is slower than neighbor joining, but is much faster than SiFit and SCITE. When the number of cells increases, the running time may increase faster than the case of increasing the number of sites for all these methods. In this case, ScisTree is still much faster than SCITE and SiFit: for example, for 500 cells and 2,500 sites, ScisTree takes three and half hours, while SiFit takes about 122 hours and SCITE takes about 42 hours. Again, running time of SCITE and SiFit depend on the parameter settings (especially the number of iterations). Note that SiFit is run with multiple threads by default. ScisTree runs with a single thread at present. Moreover, ScisTree is much faster than SCIΦ. We note that SCIΦ runs with raw sequence reads. Even excluding the time spent on reads processing, ScisTree is faster than SCIΦ by at least one order of magnitude.

**Figure 6:**
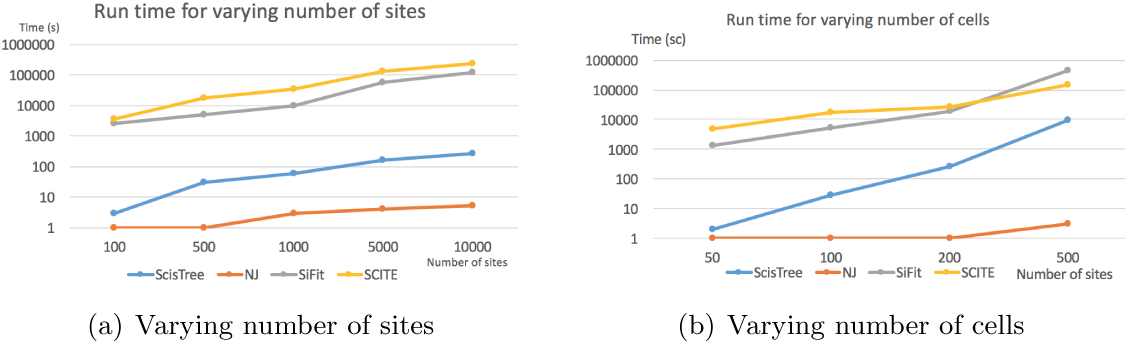
Average running time (in seconds) for a single simulated dataset of ScisTree, neighbor joining (NJ), SiFit and SCITE. Time for increasing the number of sites m (with 100 cells) is in part 6(a), and the time for increasing the number of cells n (with m being five times of n) is in part 6(b).

### 4.4 Real data

We apply ScisTree on an acute lymphoblastic leukemia dataset (Gawad *et al.*, 2014) which was analyzed by SCIΦ previously in Singer *et al.* (2018). It contains 255 cells from a patient (number 3) with acute lymphoblastic leukemia. The data is in the form of targeted DNA sequence reads from the 255 cells. We align the reads from these cells (and one additional wild-type cell used as outgroup) onto the whole human genome reference. We use Monovar to call SNV genotypes and compute the genotype probabilities. The inferred cell lineage tree is shown in the Supplemental Materials. One can see that the cell lineage tree has a side-chain shape (with 24 cells plus the wild-type cell) near the root, then diverges into two clades (one smaller and one larger). While the true tree is not known, the mutations (not shown in the tree) provide some potentially useful information. There are total 406 SNV sites (mutations) that are called by ScisTree. Among these, 53 mutations are located on the branch from the root to the clade of 255 cells. That is, these mutations are shared by all cancer cells. 173 mutations (i.e., about 43%) are located within the side-chain of 24 cancer cells, mostly along the main path from the root to the other cancer cells). This may indicate that mutations accumulate before entering the rapid growth phase of the cancer cells. In Singer *et al.* (2018), the called genotypes by SCIΦ are said to have less “noise” than the called genotypes by Monovar. ScisTree calls fewer SNV sites than Monovar: there are 808 SNV sites called by Monovar originally. However, the genotypes called by ScisTree are not as “clean” as shown in Singer *et al.* (2018). Based on our experience, SCIΦ may collapse similar clades into a single clade. While this is only a preliminary study, it shows that the inferred cell lineage tree can potentially be useful for understanding cancer evolution.

#### Additional results

Due to the space limit, some results (e.g., doublet imputation) are given in the Supplemental Materials.

## 5 Conclusion and discussions

In this paper, we present a new method for inferring cell lineage tree given individualized genotype probabilities. Individualized genotype probability allows more faithful information about single cell data and can improve inference accuracy. For example, if there is evidence for the presence of dropout at a site of a cell, we can decrease the probability of the wild-type genotype at this position for this cell to accommodate the potential dropout. We now discuss several related issues.

### Infinite sites model or not?

Whether the infinite sites model is valid for single cell evolution has not been resolved in the literature (Kuipers *et al.*, 2017; Zafar *et al.*, 2017). In this paper, we assume the IS model is valid at least for large portion of data. When the underlying evolutionary process deviates significantly from the IS model, assuming the IS model will be problematic. We note that in that case, it is difficult to distinguish technological noises (such as allelic dropouts and doublets) and complexities caused by these non-IS mutational processes in currently available data. For example, it was argued in Zafar *et al.* (2017) that the non-hereditary colorectal cancer dataset analyzed by Wu *et al.* (2016) does not agree with the IS model because the data contains many cases of violations to the IS model. However, since the dropout rate can be 20% or higher, many of the observed violations may be due to technological noises such as dropouts. In fact, the genotypes as shown in Figure 3 of Wu *et al.* (2016) appear to largely match what we expect from the IS model. For this dataset, ScisTree finds a solution where less than 7% of genotypes need to be changed to make the data fit the IS model.

### Single cell data simulation

We conduct extensive simulation on single cell sequencing in this paper. We note that single cell sequencing is complex (e.g., different amplification techniques may have different dropout effects). At present, there is no widely accepted single cell sequencing simulator. We expect a good simulator will be useful for future methodology development.

### Efficiency

ScisTree is efficient and scalable to large number of sites. Comparing with existing methods such as SCITE and SiFit, ScisTree runs much faster (often by over 100 times). Computational efficiency is likely to be a main issue for single cell analysis. With the fast developing technologies, it is expected that the size of single cell data will grow rapidly.

## Acknowledgments

I thank I. Mandoiu for useful discussions.

## Funding

This work has been supported by U.S. NSF grants IIS-1526415 and CCF-1718093.

