## Supplemental Materials for "Accurate and Efficient Cell Lineage Tree Inference from Noisy Single Cell Data: the Maximum Likelihood Perfect Phylogeny Approach"

Yufeng Wu
Department of Computer Science and Engineering
University of Connecticut
Storrs, CT 06269, U.S.A.

#### 1 Basics of the perfect phylogeny problem

Perfect phylogeny is central to the ScisTree approach. To be complete, we provide a short description for the perfect phylogeny problem. The classic perfect phylogeny problem considers a binary matrix $M$ of $n$ rows and $m$ columns. A row usually corresponds to a taxon and a column corresponds to a character (or site). For example, if $M[i,j] = 1$, this may indicate taxon $i$ has the character (say a mutation) at site $j$. Given a binary matrix $M$, a zero-rooted perfect phylogeny is a rooted phylogenetic tree where its leaves are labeled by a distinct row (taxon) in $M$, each column in $M$ labels exactly one edge (multiple columns can label the same edge), and the set of characters along the root to a leaf is exactly the set of sites where the leaf has 1 in $M$. Note that perfect phylogeny may be a multifurcating tree. It is also possible that the root is unknown. In this paper, we focus on the rooted perfect phylogeny. This is because the SNVs considered in the paper are caused by somatic mutations and therefore the root has all wild-type genotypes. Perfect phylogeny may be suitable when the rate of mutations is very low so that the chance of more than one mutation occurring at exactly the same site is small. This can be the case for DNA sequences from cells in an individual that originate from a fertilized egg cell at relatively recent time.

For two columns (sites) $a$ and $b$ and a row $h$, we call the ordered pair $M[h,a]M[h,b]$ a gamete. There are four possible gametes: 00, 01, 10 and 11. Given a binary matrix $M$, the following theorem is the main result concerning the existence of (rooted) perfect phylogeny.

**Theorem 1.1** *A binary matrix $M$ has a zero-rooted perfect phylogeny if and only if no pairs of columns $c$ and $d$ has the three gametes: $01, 10$ and $11$.*

A nice property of perfect phylogeny is that perfect phylogeny, if exists, is essentially unique. That is, when a perfect phylogeny exists, there exists a single and unique perfect phylogeny where all internal nodes have degree at least three (while the order of labels along the same branch can be arbitrary). This is a very strong property.

#### 2 Computational complexity of the maximum likelihood perfect phylogeny problem

We assume binary genotypes in this section. Recall that in the maximum likelihood perfect phylogeny (MLPP) problem, we are given uncertain genotypes for $n$ rows and $m$ columns: for each row $i$ and column $j$, the probability of being allele $g \in \{0,1\}$ is $\mathcal{P}_{g,i,j}$. We want to find a binary matrix $M$ such that $M$ allows a perfect phylogeny and also $\prod_{i=1 \ldots n, j=1 \ldots m} \mathcal{P}_{M[i,j],i,j}$ is maximized. Note that in the MLPP problem considered in this paper, we assume the root is always zero.

We now show that the rooted MLPP problem is NP complete. Consider a known NP complete problem, vertex cover (VC). In the VC problem, given a graph $G(V,E)$, we want to find the smallest number of nodes in $V$ that cover all the edges in $E$. We now reduce VC to MLPP. Given $G(V,E)$, we construct an instance of MLPP as follows. The size of the genotype matrix $M$ is $n+1$ rows by $n$ columns, where $n$ is the size of $V$. Here, $M[i,i]=0$ for each $1 \leq i \leq n$. And $M[i,j]=1$ if there is an edge between nodes $i$ and $j$. Moreover the $n+1$ row is all 1: $M[n+1,j]=1$ for all $j$. The rest of genotypes are missing. We now create the genotype probability as follows. First, for all $g=0$ in $M$ (along the main diagonal), we have $\mathcal{P}_{0,i,j}=p_m$ where $p_m=\frac{e}{1+e}$. Note that if we change to the alternative allele at these positions, the change of log-likelihood is equal to $ln(\frac{1.0-p_m}{p_m})=-1$. That is, a decrease of 1 from the maximal possible log-probability occurs when the genotype is chosen not to be the genotype as specified in $M$ at one of these positions. Also, all the positions in $M$ with genotype 1 has probability 1.0 of being 1. That is, $\mathcal{P}_{1,i,j}=1.0$ when $M[i,j]=1$. For missing values, the probability of having 0 or 1 is 0.5: $\mathcal{P}_{0,i,j}=\mathcal{P}_{1,i,j}=0.5$ when $M[i,j]$ is missing. See Figure S1 for an example on the construction. Note that changing alleles at a missing value position doesn't change the overall probability. We let the overall maximal possible log-probability of the constructed uncertain genotypes be $log\mathcal{P}_{Max}$:

$$log\mathcal{P}_{Max} = nln(p_m) + \sum_{i,j,M[i,j]\neq 0,1} ln(0.5)$$

First note there is always a solution for MLPP with log-probability no smaller than $log\mathcal{P}_{Max}-n$: we can simply change all $M[i,i]$ to 1 (with total cost n) and set all missing values to be 1 also. The resulting matrix $M'$ is all 1, which has log-probability $logP_{max}-n$.

Now we claim there is a VC of size $k$ iff there is a MLPP solution of log-probability of $log\mathcal{P}_{max}-k$. To see why this holds, consider a VC of size $k$. We then set all $M[i,i]$ to 1 for $i \in VC$. We deal with missing values as follows: for each missing value at $M[i,j]$ at row $i$ and column $j$, if $i \in VC$ or $j \in VC$, we change $M[i,j]$ to 1. Otherwise, we change $M[i,j]$ to 0. We claim this gives a valid MLPP solution with probability $log\mathcal{P}_{max}-k$. The log-probability is clearly correct: we only change $k$ main-diagonal 0s where correspond to nodes in VC, and each such change reduces the log-probability by 1.0; changes to missing value do not change the probability. We claim that there is no incompatible pairs of sites in the changed matrix $M$. To see this, note that after changing, all columns $i$ where $i \in VC$ become all-1 and thus are compatible with all other columns. Thus we only need to ensure there is no incompatibility between two columns not in VC. Let two such columns be $i$ and $j$ where $i,j \notin VC$. That is, there is no edge between nodes $i$ and $j$. So there is no 10 or 01 in the original $M$ at columns $i$ and $j$. This is because there is only a single 0 at each column $i$ in the original $M$ (at $M[i,i]$, along the main diagonal), and $M[i,j]=1$ would imply there is an edge between $i$ and $j$. We need to ensure there is no 01/10 gametes introduced in the changed $M'$. Consider a row $h$. If $h \in VC$, then the single zero in this row is changed to 1, and so are all the remaining missing values in this row. That is, there is no added zeros in row h and so there is no new 01/10 gametes in row h. Now suppose $h \notin VC$. Note to introduce 01/10 gametes, exactly one of $M[h,i]$ and $M[h,j]$ is missing (if both missing, they are both set to 0; if none is missing and so the alleles at $M[h,i]$ and $M[h,j]$ are the same as in original $M$, then it follows from the fact that there is no 01/10 in the original $M$). So let $M[h,i]$ be missing and $M[h,j]$ is not (and then $M[h,j]=1$ since we change $M[h,i]$ to 0 in this case). Now since $h$ is not in VC, and $h$ and $j$ are neighbor (since $M[h,j]=1$), $j$ must be in VC (otherwise the edge $(h,j)$ is not covered). This contradicts the assumption that none of $i$ and $j$ is in VC.

The other direction is easy: if there exists a MLPP with log-probability of $logP_{Max}-k$, then there exists a VC of size $k$. First, $k$ allele-0s are changed to 1 in $M$ because the 1-genotypes are too costly to change (changing genotype 1 to 0 will lead to negative infinite log-probability); also changing missing values doesn't change the probability. Moreover, since the changed matrix $M$ allows a perfect phylogeny, at least one 0-genotype corresponding to a node of any edge $(u, v)$ must be chosen. This is because there are 01, 10 and 11 gametes in the columns $u$ and $v$. Note that the 11 gamete is from the row $n+1$. Also 01 and 10 gametes are because of the edge $(u, v)$. Thus, at least one of the zeros must be chosen to change to 1 to allow perfect phylogeny.

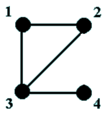


(a) A graph

| | 1 | 2 | 3 | 4 |
|---|---|---|---|---|
| 1 | 0 | 1 | 1 | * |
| 2 | 1 | 0 | 1 | * |
| 3 | 1 | 1 | 0 | 1 |
| 4 | * | * | 1 | 0 |
| 5 | 1 | 1 | 1 | 1 |

(b) Genotype matrix $M$

| | 1 | 2 | 3 | 4 |
|---|---|---|---|---|
| 1 | $\frac{e}{1+e}$ | 0.0 | 0.0 | 0.5 |
| 2 | 0.0 | $\frac{e}{1+e}$ | 0.0 | 0.5 |
| 3 | 0.0 | 0.0 | $\frac{e}{1+e}$ | 0.0 |
| 4 | 0.5 | 0.5 | 0.0 | $\frac{e}{1+e}$ |
| 5 | 0.0 | 0.0 | 0.0 | 0.0 |

(c) Genotype probability (of 0-genotypes) of $M$

| | 1 | 2 | 3 | 4 |
|---|---|---|---|---|
| 1 | 0 | 1 | 1 | **0** |
| 2 | 1 | **1** | 1 | **1** |
| 3 | 1 | 1 | **1** | 1 |
| 4 | **0** | **1** | 1 | 0 |
| 5 | 1 | 1 | 1 | 1 |

(d) Changed $M'$ allowing a perfect phylogeny

Figure S1: *Illustration of the construction for showing Vertex Cover reduces to the MLPP problem. Part 1(b) shows the constructed genotype matrix $M$ for the graph in Part 1(a). The underlying rows/columns correspond to the nodes in the vertex cover of the graph. Part 1(c) shows the corresponding genotype probabilities. Note the probabilities shown are for the probabilities of genotype 0s. Part 1(d) shows the changed matrix from $M$ that allows a perfect phylogeny where the changed 0s correspond to nodes in the vertex cover. The boldface positions are those changed from $M$ (including missing value changes).*

### 3 Additional methods

#### 3.1 Computing the maximum probability of genotypes for a fixed tree for the case of ternary genotypes

In the Methods Section, we describe an algorithm for computing the maximum probability of binary genotypes. We now consider the case of ternary genotypes, where 0 and 2 are the homozygous wild-type and mutant respectively, and 1 is the heterozygote. We follow the IS model, which imply that the genotypes 1 and 2 are caused by two different mutation events. That is, mutation $m_1$ is for the mutation in the heterozygote and mutation $m_2$ is for the second mutation causing the homozygous mutant. We make the following assumption regarding to the two mutations: $m_1$ is ancestral to $m_2$. That is, the cells carrying the $m_2$ mutations are a subset of the cells carrying $m_1$ mutations. This assumption can be justified by noting that homozygous mutant genotypes may be caused by a recurrent mutation or a loss of heterozygousity (LOH) event at a cell that carries $m_1$ mutant. Therefore, the heterzygous mutation is ancestral to the homozygous mutation. Under this assumption, we extend the maximum likelihood algorithm for binary genotypes to ternary genotypes as follows.

For each node $v$ in the given rooted binary tree $T$, we define $q_{v,g_a,g_b}$ to be the ratio of the probability of that all cells within the subtree rooted at $v$ have genotype state $g_a$ to the probability that those genotypes in this subtree have gentoype state $g_b$. Also, we define $q^m_{v,2,1} = max_{v' \in Nodes(T_v)} q_{v',2,1}$. That is, $q^m_{v,2,1}$ is the maximum of $q_{v',2,1}$ for any node $v'$ within the subtree $T_v$. Then we have the following algorithm.

**Algorithm 1** Maximum probability computation for ternary genotypes at site $s$ for a fixed binary tree $T$

**for** node $v \in Nodes(T)$ in the bottom-up order (i.e., leaves first) **do**
    **if** v is a leaf **then**
        $q_{v,1,0} \leftarrow \frac{\mathcal{P}_{1,c(v),s}}{\mathcal{P}_{0,c(v),s}}$
        $q_{v,2,1} \leftarrow \frac{\mathcal{P}_{2,c(v),s}}{\mathcal{P}_{1,c(v),s}}$
        $q^m_{v,2,1} \leftarrow q_{v,2,1}$
    **else**
        Let $v_l$ and $v_r$ being the two children of $v$.
        $q_{v,1,0} \leftarrow q_{v_l,1,0} q_{v_r,1,0}$
        $q_{v,2,1} \leftarrow q_{v_l,2,1} q_{v_r,2,1}$
        $q^m_{v,2,1} = max(q_{v,2,1}, q^m_{v_l,2,1}, q^m_{v_r,2,1})$
    **end if**
**end for**
$p_{s,G,T} \leftarrow max_{v \in Nodes(T)} max(q_{v,1,0}, q_{v,1,0} * q^m_{v,2,1}) * \prod_{c=1 \ldots n} \mathcal{P}_{0,c,s}$

The above algorithm is a dynamic programming approach which extends the algorithm for the binary genotype case. Intuitively, it stores the best branch for placing the $m_2$ mutation within the subtree $T_v$. This avoids enumerating all possible locations for the $m_2$ mutation, assuming the other mutation $m_1$ mutation is ancestral to $m_2$. The running time for the ternary genotypes is $O(n)$ for a single site, which is asymptotically the same as the binary genotype case.

#### 3.2 Doublet imputation

The key step in the doublet imputation method is scoring a candidate cell as a doublet. ScisTree takes the following approach for scoring a candidate doublet cell: ScisTree first calls genotypes of all other cells except the doublet candidate using the maximum likelihood perfect phylogeny approach; then it finds the most probable splitting of the candidate cell into two derived cells such that genotypes of the two derived cells are still consistent with the IS model; the candidate score is the maximum probability of splitting the candidate cell. Once the optimal score is obtained, genotypes can be called for the two new cells by optimally splitting a doublet cell into two derived cells. We have a polynomial time algorithm for computing the doublet score for a candidate cell given genotypes that are called from other cells. The algorithm is given in the next section, which is a little complex.

##### 3.2.1 A polynomial time algorithm for scoring a candidate doublet cell

This section needs basic properties for perfect phylogeny as outlined in Section 1 in the Supplemental Materials. For simplicity, we assume the genotypes are binary. We are given a particular cell $c$ that is the candidate for a doublet. Here is the high-level idea on computing a doublet score for $c$. We first call the genotypes of the remaining cells (i.e., discarding $c$ from the genotypes) using the approach described in the main text of this paper. The called genotypes $G_c$ satisfy the IS model. Let two new cells $c_1$ and $c_2$ be the ones forming the doublet $c$. The objective is finding binary genotypes $g_1$ and $g_2$ of length $m$ for $c_1$ and $c_2$, such that the combined genotype $g = g_1 \cup g_2$ has the highest probability regarding to the doublet cell $c$, and adding $g_1$ and $g_2$ to $G_c$ still satisfies the IS model. Note that $c_1$ and $c_2$ may or may not be present in the original list of cells (so that we cannot just simply try to combine pairs of existing cells to form the doublet). Since there are $m$

sites, there are four ordered pairs of genotypes for $c_1$ and $c_2$ at each site: 0/0, 0/1, 1/0 and 1/1. When $m$ is large, we cannot enumerate all the combinations of choices for the $m$ sites. We now present an efficient algorithm for calculating the maximum score and finding the optimal doublet genotypes for a candidate cell.

Our algorithm is based on the clade tree $T_c$, which is related to the cell lineage tree $T$. Recall that $G_c$ satisfies the IS model and so the clades in $G_c$ are nested in a tree (the clade tree). The IS model implies the so-called "clade property": the clades are compatible with the IS model iff any two clades are either disjoint or one contains the other. The nodes (except the root) in $T_c$ are the clades implied by sites in $G_c$. The root of $T_c$ has the clade containing all cells in $G_c$. In the following, we use site, clade and node interchangeably when no confusion is introduced. For each node $v$, the children of $v$ are those clades contained by $v$ and are maximal. That is, if a clade $v_1$ is properly contained in $v_2$, and $v_2$ is properly contained in $v_3$. Then $v_1$ is not a child of $v_3$. Duplicate clades are assumed to be absent from $T_c$ [1]. We need to determine the genotypes at each site for the new cells $c_1$ and $c_2$. That is, if the new genotype is 1 at a new cell say $c_1$ at a node (site) $v$, $c_1$ is inserted into the clade at $v$. The key observation is that to ensure the clade property holds after adding $c_1/c_2$, we only need to ensure the clades at a node and each of its descendants are compatible with the IS model when the two new cells are added. This allows a dynamic programming algorithm on clades in a bottom-up order. Note that when the two new cells are added, we need to determine their genotypes at each site (clade). In this algorithm, we consider each clade (site): adding the two new cells must make the clades remain consistent with the IS model. Now we present more details.

We first note that if two clades are disjoint before adding $c_1/c_2$, they remain disjoint. This is because these two clades cannot have one contain another because we can only decide whether to add $c_1/c_2$ to these two clades, not the other cells; since the cells are disjoint before, there are cells that are private to both cells; thus they cannot have containment relationship. Similarly, we can see that if two clades have one contain another, then they must remain so after adding $c_1/c_2$. This implies that the clade tree remains the same topology after inserting $c_1/c_2$.

For each node $v$, we define $m_v(h_1, h_2)$ as the maximum probability such that the new genotypes $c_1$ and $c_2$ are contained by the clade at $v$, the new genotype of $c_1/c_2$ at site $v$ has genotype states $h_1/h_2$ and the genotypes of the entire subtree satisfy the IS model. There are four choices for $h_1/h_2$: 0/0, 0/1, 1/0, and 1/1. Then the overall maximum probability is equal to $m_r(1, 1)$, where $r$ is the root. This is because the root clade contains all cells (and so we have 1 and 1 for both $c_1$ and $c_2$ at the root). At a leaf clade $v$, $m_v(0, 0) = \mathcal{P}_{0,c,v}$, and $m_v(0, 1) = m_v(1, 0) = m_v(1, 1) = \mathcal{P}_{1,c,v}$. Now consider an internal clade $v$. We have the following cases.

**Case 1**: for $m_v(0, 0)$ where both $c_1$ and $c_2$ have genotype 0 at site $v$. We claim in this case, the subtree rooted at $v$ agrees with the IS model iff all children nodes $v_1$ below $v$ have genotypes 0/0 for $c_1/c_2$. To see this, we first argue that if the subtree at $v$ agrees with the IS model, all children of $v$ have 0/0 genotypes at $c_1/c_2$. This is because the clade of allele 1s of a site (node) $v_1$ below $v$ is strictly contained by the clade of the site $v$. Note that $c_1/c_2$ having genotypes 0/0 implies none of $c_1$ and $c_2$ is inserted to the clade at $v$. Thus, none of its children has either $c_1$ or $c_2$ in its clade too (and so each child has genotype 0/0 in $c_1/c_2$). On the other hand, if all children of $v$ have genotypes 0/0 at $c_1/c_2$, then only gametes 00 are present at cells $c_1$ and $c_2$, which don't lead to violation of the IS model. Thus,

[1]If duplicate clades are present, we may introduce weights to the clades. That is, we assign the weight of clade to be 2 if the clade appears twice. The algorithm can be easily modified to consider weights.

$$m_v(0,0) = \mathcal{P}_{0,c,v} + \sum_{v' \in Child(v)} m_{v'}(0,0)$$

**Case 2**: for $m_v(0,1)$ ($m_v(1,0)$ is the similar). Here, $c_1$ has genotype 0 and $c_2$ have genotype 1 at site $v$. Then we claim in this case, the subtree rooted at $v$ agrees with the IS model iff either at each child of $v$ the genotypes are 0/0 at $c_1/c_2$ or or at exactly one child of $v$ has 0/1 at $c_1/c_2$ and the rest are 0/0. To see this, first note that all children of $v$ have genotype 0 at $c_1$ (otherwise the clade at $v$ cannot contain the clade at a child after inserting $c_1/c_2$). Suppose all children have 0/0 genotypes. Then none of $c_1$ and $c_2$ is inserted into the clades of the children of $v$, and thus they remain disjoint and also are contained by the clade at $v$ (and thus the clade property still holds). The same is true when at most one has a 0/1 genotypes at $c_1/c_2$. Now suppose there are at least two children with 1 genotypes at $c_2$. Then the clades at these two children won't be disjoint (and has no containment relationship either). This leads to violation of the clade property. So,

$$m_v(0,1) = \mathcal{P}_{1,c,v} + min(\sum_{v' \in Child(v)} m_{v'}(0,0), min_{v_i \in Child(v)}(m_{v_i}(0,1) + \sum_{v' \in Child(v), v' \neq v_i} m_{v'}(0,0)))$$

Here $Child(v)$ is the set of nodes that are children (i.e., direct descendants) of $v$.

**Case 3**: for $m_v(1,1)$ where both $c_1$ and $c_2$ have genotype 1 at site $v$. Then we claim that in this case, the subtree rooted at $v$ agrees with the IS model iff one of the following holds: (i) at all children of $v$ $c_1/c_2$ have genotypes 0/0, (ii) at exactly one site $v'$ as a child of $v$ new genotypes of $c_1/c_2$ are 0/1 or 1/0 or 1/1 and all other genotypes are 0/0, or (iii) at one child node (site) the new genotypes of $c_1/c_2$ are 0/1 and another child site has new genotypes 1/0 and all other children have new genotypes 0/0. The cases (i) and (ii) are similar to the previous two cases. The case (iii) can be seen by noting that if there are more than one child node having say 0/1 genotypes, this leads to exactly the same situation as in Case 2 (which violates the clade property). So,

$$\begin{aligned} m_v(1,1) = \mathcal{P}_{1,c,v} + min(&\sum_{v' \in Child(v)} m_{v'}(0,0), min_{v_i \in Child(v)}(min(m_{v_i}(0,1), m_{v_i}(1,0), m_{v_i}(1,1)) \\ &+ \sum_{v' \neq v_i} m_{v'}(0,0)), min_{v_i, v_j \in Child(v), v_i \neq v_j}(m_{v_i}(1,0) + m_{v_j}(0,1) + \sum_{v' \in Child(v), v' \neq v_i, v_j} m_{v'}(0,0))) \end{aligned}$$

Now in all the above cases, when new cells $c_1$ and $c_2$ are added, the clades in the clade tree remain either disjoint or one contains another. This is because each of the new cells is only added along one child of a clade; thus, two children of the clade remain disjoint after adding new cells. Moreover, when a new cell is added to a clade, then its parent must have also added this new cell. This way, if a clade contains another clade, then it remains so after new cells are added. Thus, when the new genotypes found by the algorithm are added to the clades, the clades are still compatible with the IS model. The optimal genotypes for the two new cells can be found by traceback.

**Time analysis for doublet imputation**. Recall that there are $n$ cells and $m$ sites. The number of clades in the clade tree is $O(m)$. The dynamic programming algorithm over the clade tree can be implemented to run $O(m^2+mn)$. Let the number of doublets be $n_d$. Finding each of $n_d$ doublets, we need to consider $O(n)$ doublet candidates. In each iteration, we need to consider $O(n)$ neighboring trees found by nearest neighbor interchange (NNI). Thus the doublet imputation algorithm runs in $O(n_d n^2 m(m+n)N) = O(n^3 m(m+n)N)$ time. Here, $N$ is the number of iterations for convergence for finding the maximum likelihood tree.

### 4 Simulation

#### 4.1 Simulation of a true cell lineage tree

We first generate a random tree topology with $n$ taxa. We then assign branch lengths to the tree. There are different ways to choose branch lengths, which depend on the underlying evolutionary model of single cells. One simple approach is assuming the cell lineage tree satisfies the clock property. However, enforcing the strict clock property may not be realistic in real data since mutation rates along different branches may vary. We choose the following way of assigning branch lengths. First, we assign randomly chosen initial branch lengths which fit the clock model. We normalize the total tree height to be unit height. Note that only the relative magnitude (not the absolute magnitude) of different branch lengths matter because we fix the number of SNV sites in our simulation. Then, to model the deviation from the clock property, we add a random perturbation (in the range from 0.0 to $t_0$) to each branch length. By default, $t_0 = 0.1$ (i.e., the maximum perturbation is 10% of the tree height). The resulting cell lineage tree with branch lengths is used as the true cell lineage tree $T_{sim}$.

#### 4.2 Mutation simulation on a given cell lineage tree

Given a cell lineage tree $T_{sim}$ (with branch lengths), we simulate $m$ SNV sites by simulating mutations on $T_{sim}$. The value of $m$ is provided by the user. By default, mutations follow the IS model. Each mutation is placed on a stochastically chosen branch. The probability of a branch being chosen is proportional to the length of the branch.

In practice, more complex evolutionary processes may occur. These include recurrent mutations and copy number aberrations (gain or loss). We allow a fraction $r_m$ of sites to have recurrent mutations. For each of these sites, we simulate one additional mutation at the same site. $r_m$ is between 0 and 1, and usually small. In this simulation, similar to most existing literatures, we don't consider the situation of the copy number gain. We do allow copy number loss events. Copy number loss events occur in the rate $m_c$ per unit length. Usually $m_c$ is a small value. When copy number loss occurs at a site, we drop one arbitrary allele from the current cell at the site where the event occurs.

#### 4.3 Genotype simulation

After mutations are simulated, true genotypes are decided for all cells. A true genotype of a cell at a site can be represented by a pair of integers $(n_0, n_1)$. Here $n_0$ (respectively $n_1$) is the number of wild-type (respectively mutant) alleles. Note that if there is no copy number change, $n_0 + n_1 = 2$. We then simulate the noisy genotypes by imposing technological errors during the process of obtaining observed genotypes from true genotypes. This involves dropouts, which occur at rate $\epsilon_d$. It also involves genotyping errors, which occur at rate $\epsilon_g$. Usually $\epsilon_d$ can be up to 20% or higher, which is much larger than $\epsilon_g$ in the current technologies. Binary genotypes are formed by considering genotypes with no mutants being genotype 0. and the others being genotype 1. Ternary genotypes can be formed by considering the homozygotes and heterozygotes as usual. If the simulated genotype has only one allele (due to copy number loss), it is considered to be a homozygote. Genotype probability can be obtained as follows. By default, a simulated genotype 0 has probability $1.0 - FN$, where false negative rate $FN \approx \epsilon_d + \epsilon_g$. A simulated genotype 1 has probability $1.0 - FP$, where false positive rate $FP \approx \epsilon_g$. This is because without other information, a simulated genotype 0 may be caused by a dropout from a true genotype 1 or due to a genotyping error; a simulated genotype 1 may be a result of genotyping error from a true genotype 0. This setting is called the uniform genotype probability. Note that although more complex probability calculation can be done, we choose this simple scheme because in reality parameters are not known; using such crude probability calculation may reveal how well various inference methods can work with noisy input.

Doublets can be simulated by randomly pairing genotypes from two cells. The new doublet cell is added to the data, while the two source cells are removed.

### 5 Additional results

#### 5.1 Doublet imputation on simulated data

Current single cell data may contain doublets. When processing such single cell data, one may try to integrate doublets into inference. For example, one may infer phylogeny with doublets. In this paper, we take a slightly different approach for dealing with doublets. Instead of considering doublets in specific applications, we aim to imputing doublets using the approach described in Section 3.2. That is, we want to find which cells are doublets. And for doublet cells, we want to impute their genotypes of the individual cells forming the doublets. This way, we may replace genotypes from doublet cells with imputed genotypes from individual cells.

We simulate single cell genotypes with doublets as follows. We use the uniform genotype probability model here. We fix $n$ to be 100. The default value of $m$ is $1,000$. The default number of doublets is 5 (i.e., about 5% doublet rate). The rest of parameters are in the main text of this paper. We assume the number of doublets is known when running ScisTree. We use two statistics for measuring the performance of doublet imputation. (i) Accuracy of doublet calling: percentage of true doublets called. For example, if there are two simulated doublets in the data and ScisTree finds one correctly among the two called doublets. Then the accuracy is 50%. (ii) For correctly called doublets, the accuracy of imputed genotypes of individual cells forming the doublets. Suppose a doublet is called correctly. ScisTree outputs the imputed genotypes of the two cells forming this doublet. The accuracy of called genotypes for a true doublet is equal to the average percentage of correctly called genotypes in these two individual cells. Figure S2 shows the results of doublet imputation. We can see that doublet imputation works reasonably well in the simulations. ScistTree is able to impute about 80% doublets when there is a single doublet in 100 cells. Note that random guess is expected to have accuracy of 1% in this case. When dropout rate is low (say at 1%) or the number of sites is large (say $2,000$), double imputation accuracy can be around 80%. Even when the doublet percentage is relatively high say at 20%, we can still impute more than 50% doublets correctly. For correctly called doublets, imputed genotype accuracy is high (usually above 95%). The main case where doublet imputation doesn't perform well is when the number of sites is small (say $m = 100$). This is expected because when the number of sites is small, evidence of doublets is weak in the data. Our results show that doublet imputation works reasonably well.

#### 5.2 Inference results on the acute lymphoblastic leukemia dataset

Figure S3 shows the cell lineage tree from 255 acute lymphoblastic leukemia cells and a wild-type cell (used as outgroup to root the tree). There are 406 SNV sites (not shown) called by ScisTree.

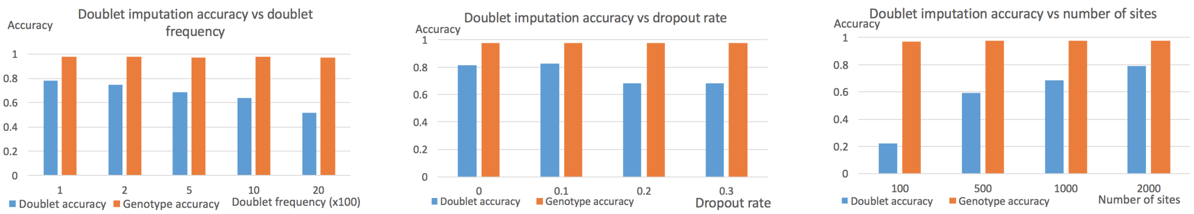


(a) Varying doublet frequency (b) Varying dropout rates (c) Varying number of sites

Figure S2: *Doublet imputation by ScisTree. Doublet accuracy: percentage of correctly imputed doublet cells. Genotype accuracy: fraction of correctly imputed genotypes of two individual cells for correctly imputed doublet cells. Part 2(a): increase the doublet frequency from 1% to 20%. Part 2(b): increase dropout rates. Part 2(c): increase the number of sites.*

#### 5.3 Cell lineage tree inference for data with doublets

Doublet is a common source of noise in single cell data and may lead to poor downstream analysis. To evaluate the impact of doublets on cell lineage tree inference, we simulate single cell data with doublets as follows. We simulate 50 cells on 500 SNV sites. We simulate various number of doublets, from 1 to 10. Then we compare the cell lineage tree inference accuracy for:

1. Impute: inference of trees on data with imputation (i.e., imputing doublets first and then performing cell tree inference on imputed data)

2. No impute: inference of trees on data without imputation (i.e., directly infer cell tree on data with doublets).

The results are shown in Figure S4. The inference error is the average Robinson-Foulds distance among 50 replicates. We can see that imputing doublets does improve the cell lineage tree accuracy. As expected, when the number of doublets is small, the difference between the two cases is not very large. Note that since the above two cases (impute or no impute) can lead to different set of taxa (since doublet imputation leads to more cell lineages), we only use non-doublet cells in comparison (i.e., if a cell is imputed to be a doublet, ignore this cell in the tree comparison) of both cases. This leads to the same set of cell lineages in comparison.

#### 5.4 Cell tree inference with copy number changes and recurrent mutations on simulated data

We now show tree inference results with two types of complexities: recurrent mutations and copy number losses. Here, we assume uniform genotype probability. Results are shown in Figure S5. It can be seen that ScisTree and SCITE have similar accuracy and perform the best overall, with SCITE being slightly more accurate in some cases. ScisTree and SCITE consistently outperform neighbor joining in many settings. SiFit appears to be less accurate in our tests.

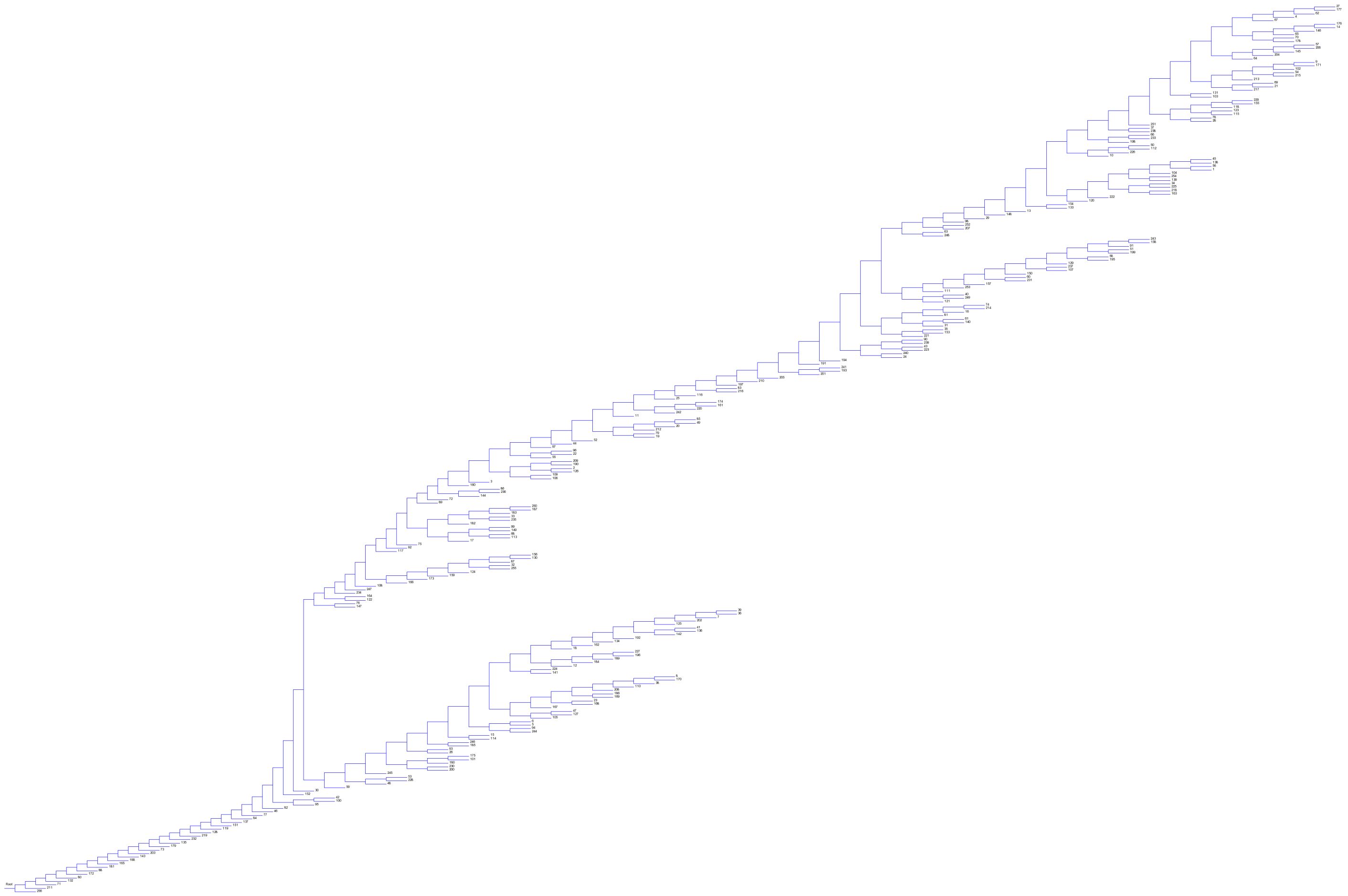

Figure S3: *Inferred cell lineage tree from an acute lymphoblastic leukemia dataset from the patient 3. There are 255 cancer cell (numbered from 1 to 255) plus a wild-type cell (numbered 256).*

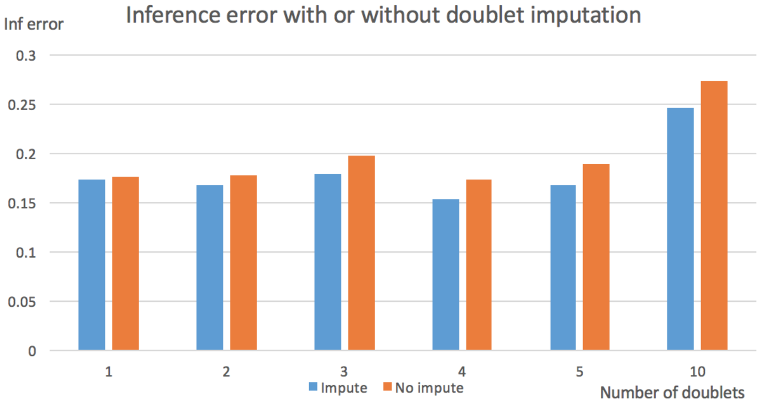


Figure S4: *Cell lineage tree inference error with doublets. 50 cells and 500 SNV sites. Impute: tree inference error with data where doublets are first imputed. No impute: tree inference error with data where no doublet imputation is performed. Number of simulated doublets: 1, 2, 3, 4, 5 and 10. Inference error is the average RF distance over 50 replicates.*

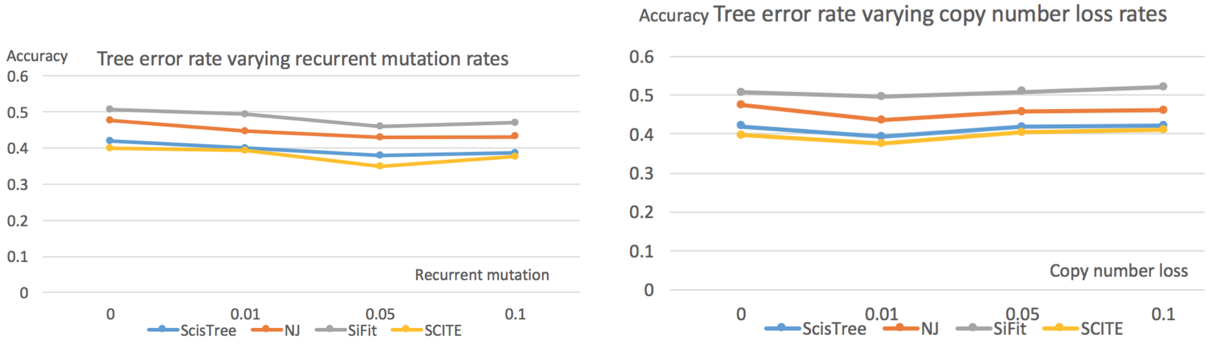


(a) Varying recurrent mutation rates

(b) Varying copy number loss rate

Figure S5: *Cell lineage tree inference error with copy number changes and recurrent mutations by ScisTree. Part 5(a) shows the effect of recurrent mutation error rate. Part 5(b) shows the effect of copy number loss rate. Y axis: average RF distance between true tree and inferred tree. Default settings of parameters are used for parameters not being tested.*
